# Homeostasis of protein and mRNA concentrations in growing cells

**DOI:** 10.1101/255950

**Authors:** Jie Lin, Ariel Amir

## Abstract

Many experiments show that the numbers of mRNA and protein are proportional to the cell volume in growing cells. However, models of stochastic gene expression often assume constant transcription rate per gene and constant translation rate per mRNA, which are incompatible with these experiments. Here, we construct a minimal gene expression model to fill this gap. Assuming ribosomes and RNA polymerases are limiting in gene expression, we find that (1) because the ribosomes translate all proteins, the concentrations of proteins and mRNAs are regulated in an exponentially growing cell volume; (2) the competition between genes for the RNA polymerases makes the transcription rate independent of the genome number. Furthermore, by extending the model to situations in which DNA (mRNA) can be saturated by RNA polymerases (ribosomes) and becomes limiting, we predict a transition from exponential to linear growth of cell volume as the protein-to-DNA ratio increases.

---

Despite the noisy nature of gene expression [1–6], various aspects of single cell dynamics, such as volume growth, are effectively deterministic. Recent single-cell measurements show that the growth of cell volumes is often exponential. These include bacteria [7–10], archaea [11], budding yeast [10, 12–15] and mammalian cells [10, 16]. Moreover, the mRNA and protein numbers ı are often proportional to the cell volume throughout the cell cycle: the homeostasis of mRNA concentration and protein concentration is maintained in an exponentially growing cell volume with variable genome copy number [17–22]. The exponential growths of mRNA and protein number indicate dynamical transcription and translation rates proportional to the cell volume, and also independent of the genome copy number. However, current gene expression models often assume constant transcription o rate per gene and constant translation rate per mRNA ı (constant rate model) [1, 5, 23–25]. Assuming a finite degradation rate of mRNAs and non-degradable proteins, these models lead to a constant mRNA number proportional to the gene copy number and linear growth of protein number [26–28], incompatible with the proportionality of mRNA and protein number to the exponentially growing cell volume.

Since the cell volume, protein copy number and mRNA copy number grow exponentially throughout the cell cycle, one may expect a sufficient condition to achieve a constant concentration is to let them grow with the same exponential growth rate. However, mathematical analysis suggests this is insufficient. Let us consider the logarithm of protein concentration *c*, which can be written as ln(*c*) = ln(*p*)-ln(*V*). Here *p* is the protein number and *V* is the cell volume. If one assumes the protein number and the cell volume grow exponentially but independently, with time-dependent exponential growth rates λ_*p*_(*t*) and λ_*ν*_(*t*) respectively, the time derivative of the logarithm of concentration then obeys *d*ln(*c*)/*dt* ~ λ_*p*_(*t*)–λ_*ν*_(*t*). Even when the time-averaged growth rates of protein number and cell volume are equal, 〈λ_*p*_(*t*)〉 = 〈λ_*ν*_(*t*)〉, any fluctuations in the difference between them will accumulate and lead to a random walk behavior of the logarithm of concentration. The homeostasis of protein and mRNA concentrations implies that there must be a regulatory mechanism in place to prevent the accumulation of noise over time.

The main goal of this work is to identify such a mechanism by developing a coarse-grained model taking into account cell volume growth explicitly. Specially, we only consider continuously proliferating cells and do not take account of non-growing cells, e.g., bacterial cells in stationary phase [29]. The ubiquity of homeostasis suggests that the global machinery of gene expression, RNA polymerases (RNAPs) and ribosomes, should play a central role within the model. Based on the assumption that the number of ribosomes is the limiting factor in translation, we find that the exponential growth of cell volume, protein number originates from the auto-catalytic nature of ribosomes [30–32]. The fact that ribosomes make all proteins ensures that the protein concentrations do not diverge. Based on the assumption that the number of RNAP is the limiting factor in transcription, we find that the mRNA number also grows exponentially and the mRNA concentration is independent of the genome copy number because of the competition between genes for this global resource [18–20]. We also study the effects of genome replication. Due to the heterogeneous timing of gene replication, the transcription rate of one gene has a cell cycle dependence. Within our model, it doubles immediately after the gene is replicated and decreases gradually as other genes are replicated. Finally, we extend our model to more general situations in which an excess of RNAP (ribosome) leads to the saturation of DNA (mRNA). We propose a phase diagram of gene expression and cellular growth controlled by the protein-to-DNA ratio. We predict a transition from exponential growth to linear growth of cell volume as the protein-to-DNA ratio passes a threshold.

## RESULTS

### Model of stochastic gene expression

In constant rate models, the transcription rate per gene and the translation rate per mRNA are constant [1, 5, 24] (Figure 1a). This implies that the gene (mRNA) number is the limiting factor in transcription (translation). Constant rate models predict a constant mRNA number proportional to the gene copy number and independent of the cell volume. However, experimental observations on plant and mammalian cells have revealed a proportionality between mRNA number and cell volume for cells with a constant genome copy number [18–20]. Moreover, even comparing the cells before and after the genome replication (S phase), the proportionality coefficient between mRNA and cell volume does not exhibit any obvious change. In contrast, a constant transcription rate per gene would predict a doubled transcription rate after the replication of the whole genome, leading to a higher mRNA concentration. In one class of constant rate models [26, 27, 33], a deterministic exponential growth of cell volume is explicitly considered. The resulting perturbation on the concentrations due to genome replication is suppressed in the long lifetime limit, but still significant for short lifetime molecules, e.g., mRNA (see Fig.1 in Ref. [27]).

**Figure 1.**
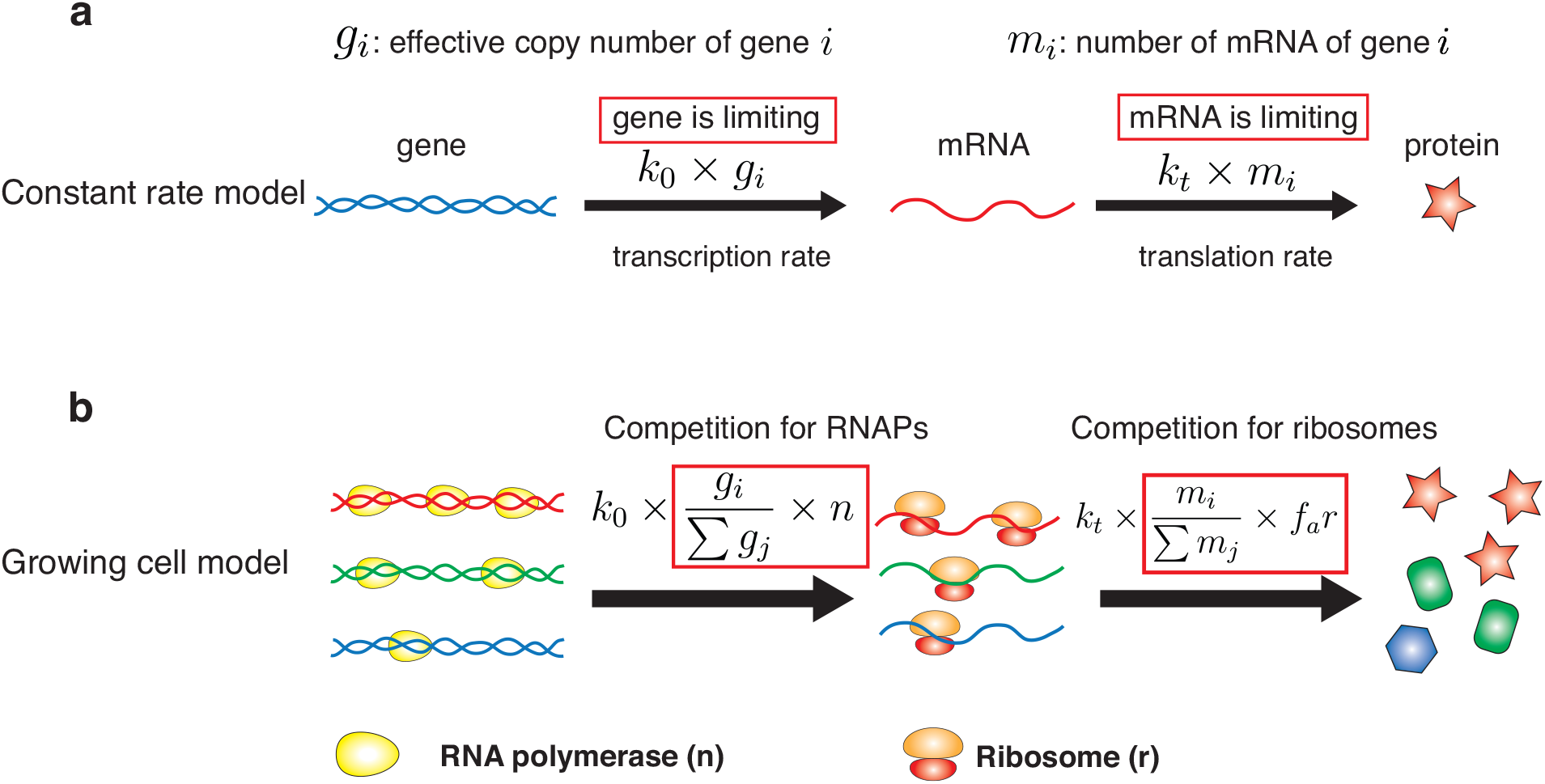
The growing cell model of stochastic gene expression in comparison with constant rate models. (a) In the constant rate model, the transcription rate is proportional to the gene copy number, and the translation rate is proportional to the mRNA number. These assumptions imply that the gene number and mRNA number are the limiting factors in gene expression. (b) In Phase 1 of the growing cell model, we introduce as limiting factors RNA polymerases (RNAPs) and ribosomes. Genes with different colors are transcribed with different rates. Here *k*_0_ is a constant and the gene regulation is coarse-grained into the gene allocation fraction *ϕ_i_* = *g_i_*/Σ_*j*_ *g_i_*. *g_i_* is the effective copy number of gene *i* (also accounting for the promoter strength). *n* is the total number of RNAPs. Translation rates of mRNA depend on the number of active ribosomes (*f_a_r*), the translation rate *k_t_*, and the fraction of mRNA i in the total pool of mRNA. In a later section (A unified phase diagram of gene expression and cellular growth), we will relax our assumptions and consider situations in which the limiting factors of gene expression become the gene number and the mRNA number.

Considering translation, various experiments have shown that the number of ribosomes is the limiting factor rather than the number of mRNAs. The most direct evidence is the growth law: the growth rate of cells is proportional to the fraction of ribosomal proteins in the total proteome (with a constant factor depending on the growth condition) [34] both for bacterial cells [30, 35] and budding yeast cells [31]. This means a constant fraction of ribosomes are actively translating mRNAs. These results suggest that in general cells are below the saturation limit in which there are too many ribosomes that the mRNAs can bind. We will therefore assume the biological situation in which mRNAs in the cell compete for the limiting resource of actively working ribosomes, therefore the translation rate of one type of mRNA is proportional to the number of active ribosomes times its fraction in the total pool of mRNAs.

Considering transcription, experiments have shown that mutants of fission yeasts altered in cell size regulated global transcription to maintain similar transcription rates per cell volume regardless of the cellular DNA content. The changes in total transcription correlated with coordinated changes in gene occupancy by RNA polymerases [36]. These results suggest that the number of RNAPs may be the limiting factor in transcription rather than the gene number, and similar evidence has been shown for bacterial cells [37] and mammalian cells [38]. However, in the same experiments on fission yeast [36], it has also been found that in cell-cycle-arrested mutants, total transcription rates stopped increasing as the cell volume exceeded a certain value, which suggested DNA became limiting for transcription at low DNA concentration. This result suggests that an excess of RNAPs may lead the gene number to become the limiting factor in certain conditions. In this section, we will focus on the scenario that both RNAP and ribosome are limiting in gene expression, which we denote as Phase 1. In this phase, we will show that the mRNA number and the protein number are proportional to the cell volume and grow exponentially. In a later section (A unified phase diagram of gene expression and cellular growth), we will consider a more general model in which the limiting nature of RNAPs and ribosomes may break down and the dynamics of mRNA and protein number is different.

To address the limiting nature of RNAP, we define an effective gene copy number *g_i_* for each gene to account for its copy number and the binding strength of its promoter, which determines its ability to compete for RNAPs. The transcription rate for one specific gene *i* is proportional to the fraction of RNAPs that are working on its gene(s), *ϕ_i_* = *g_i_*/Σ_*j*_ *g_j_*, which we denote as the gene allocation fraction. Gene regulation is thus coarse-grained into the gene allocation fraction *ϕ_i_*. The transcription rate is independent of the genome copy number since a change in the genome number leaves the allocation fraction of one gene invariant, a conclusion which is consistent with a number of experimental results on various organisms [18–20, 36].

In fact, explicit gene regulation can also be included in our model (Methods), with a time-dependent *g_i_*. In such scenarios, *g_i_* may be a function of protein concentrations (for instance, the action of transcription factors modifies the transcription rate). Such models will lead to more complex dynamics of mRNA and protein concentrations. However, since we are interested in the global behavior of gene expression and cell volume growth, we do not focus on these complex regulations in this manuscript. Our conclusions regarding the exponential growth of mRNA and protein number for constitutively expressed genes and the exponential growth of cell volume on the global level are not affected by the dynamics of gene expression of particular genes.

In the following, *m, p, r, n* represents the numbers of mRNA, protein, ribosome and RNA polymerase, respectively. Proteins (*p*) also include RNAPs (*n*) and ribo somes (*r*) [30]. We consider the degradation of mRNA with degradation time τ for all genes. The protein number decreases only through cell divisions (though adding a finite degradation rate for proteins does not affect our results). The chemical reactions of gene expression within Phase 1 of our model are summarized in the following sets of equations and Figure 1b,

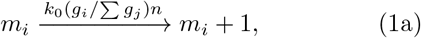

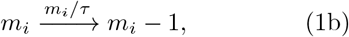

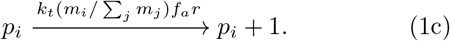

Here *k*_0_, *k_t_* are constants, characterizing the transcription (translation) rate of a single RNAP (ribosome). *f_a_* is the fraction of active ribosomes. For simplicity, we first assume the values of *ϕ_i_* do not change in time. This can be formally thought of as corresponding to an instantaneous replication of the genome. Because within our model, the transcription rate is proportional to the relative fraction of one gene in the total genome rather than its absolute copy number, the invariance of mRNA and protein concentrations before and after the genome replication is a natural result and does not rely on a long lifetime of the molecule under consideration in contrast to the constant rate models [27].

In reality, a finite duration of DNA replication and the varying time of replication initiation for different genes lead to *ϕ_i_*’s that change during the DNA replication. We later analyze a more complete version of the model which includes these gene dosage effects, but we first consider the simplified scenario of constant *ϕ_i_* that will capture the essential features of the problem. We assume the cell volume is approximately proportional to the total protein mass, i.e., *V ∝ M* = Σ_*j*_ *P_j_*, which is a reasonable approximation for bacteria [39, 40] and mammalian cells [17]. To simplify the following formulas, we consider each protein has the same mass and set the cell density as 1.

Due to the fast degradation of mRNA compared with the cell cycle duration [41, 42], the mRNA number can be well approximated as being in steady state. We can express the ensemble-averaged number of mRNA of gene *i* as

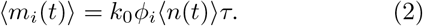

Equation (1c) then leads to the time-dependence of average ribosome number, *d*〈*r*〉/*dt* = *k_t_f_a_ϕ_r_*〈*r*〉, reproducing the auto-catalytic nature of ribosome production and the growth rate

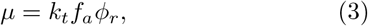

determined by the relative abundance of active ribosomes in the proteome [30, 31].

Similarly, we can find the number of protein i grows as *d*〈*p_i_*〉/*dt* = *k_t_f_a_ϕ_i_*〈*r*〉. As the cell grows and divides, the dynamics becomes insensitive to the initial conditions, so the protein number will grow exponentially as well [21]. The ratio between the averages of two protein numbers in the steady state is set by the ratio of their production rate, therefore 〈*p_i_*〉/〈*p_j_*〉 = *ϕ_i_*/*ϕ_j_*. The average number of mRNA traces the number of RNA polymerases according to Equation (2), and therefore also grows exponentially.

Throughout the cell cycle we have

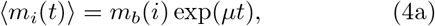

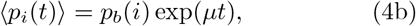

where *m_b_*(*i*) (*p_b_*(*i*)) is the number of mRNA (protein) of gene *i* at cell birth.

The concentration of mRNA and protein of gene *i* as *cm_i_ = m_i_/V, c_i_ = p_i_/V*. According to Equations (1a–1c), the deterministic equations of the above variables become (see details in Methods)

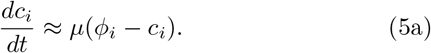

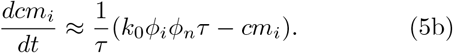

Stable fixed points exist for the dynamics of *c_i_* and *cm_i_*, which are *ϕ_i_* and *k*_0_*ϕ_i_ϕ_n_τ*. The stability of the fixed points is due to the global nature of RNAPs and ribosomes: any noises arising from the copy number of RNAPs (ribosomes) equally affect all mRNAs (proteins), and therefore leave the relative fraction of one type of mRNA (protein) in the total pool of mRNAs (proteins) invariant. The average concentrations of mRNA and protein of gene *i* become 〈*c_i_*〉 = *ϕ_i_*, and 〈*cm_i_*〉 = *k*_0_*τϕ_i_ϕ_n_*. The results are independent of the cell volume and genome copy number agreeing with experimental data on various organisms [18–20, 22].

If we only consider the dynamics of concentrations, we do not need to introduce cell division. However, taking account of cell division is necessary for the study of protein number dynamics at the single-cell level, and also important for the study of effects of gene replication. Therefore, we take explicitly cell division into account and, for concreteness, use the “adder” model for cell division by considering an initiator protein *I*. We consider an initiator protein *I*, which accumulates from 0 after cell birth, and triggers the cell division once *I* reaches the division threshold *I_c_* and is then destroyed (or “reset”, e.g., after initiation of DNA replication in bacteria, the ATP-bound DnaA is dephosphorylated to the ADP-bound form) [43–45]. During a division event, we assume proteins and mRNAs are divided between the two daughter cells following a binomial distribution [46]. The initiator protein sets the scale of absolute protein number, and the average number of proteins produced in one cell cycle is equal to Δ(*i*) = *I_c_ϕ_i_/ϕ_I_* [44]. Since the protein number grows twofold during one cell cycle, the average protein number of gene *i* at cell birth is *p_b_*(*i*) = *I_c_ϕ_i_/ϕ_i_* and the corresponding average mRNA number at cell birth is *m_b_*(*i*) = *k*_0_*I_c_τϕ_i_ϕ_n_/ϕı*. We remark that the exact molecular mechanism of cell division does not affect our results.

We corroborate the above analytical calculations with numerical simulations. These will also capture the stochastic fluctuations in gene expression levels, which are not included in the previous analysis. Due to the short lifetime of mRNAs, the production of proteins can be approximated by instantaneous bursts [24]. We introduce the burst size parameter *b*_0_ as the average number of proteins made per burst, *b*_0_ = *k_t_f_a_*〈*r*(*t*)〉/〈Σ*_j_ mj*〉 × *τ* ~ *k_t_f_a_ϕ_r_*/(*k*_0_*ϕ_n_*), independent of the cell volume. *ϕ_i_* for *N* = 200 proteins are uniformly sampled in logarithmic space, with the sum over *ϕ_i_* (including ribosome and RNAP) constraint to be precisely one. We choose the parameters to be biologically relevant for bacteria: the doubling time *T* = ln(2)/*μ* = 150 min, *r_b_* = 10^4^, *n_b_* = 10^3^, *b*_0_ = 0.8, *I_c_* = 20, *ϕ_r_* = 0.2, *f_a_* = 0.7 and *τ* = 3.5 min, see other numerical details in Methods. Our conclusions are independent of the specific choice of parameters.

In Figure 2a, we show the typical trajectories from our simulations of cell volume, protein number and mRNA number for the same gene over multiple generations. To verify the exponential growth of protein and mRNA, we average the protein and mRNA numbers given a fixed relative phase in the cell cycle progression, which is normalized by the generation time and changes from 0 to 1. The averaged values of protein and mRNA numbers (circles) are well predicted by exponential growth, Equations (4a, 4b) (black lines) without fitting parameters, as shown in Figure 2b with 3 single trajectories in the background. We also simulate a regulated gene with a time-dependent gene copy number and obtain qualitatively similar results (Methods, Figure S1).

**Figure 2.**
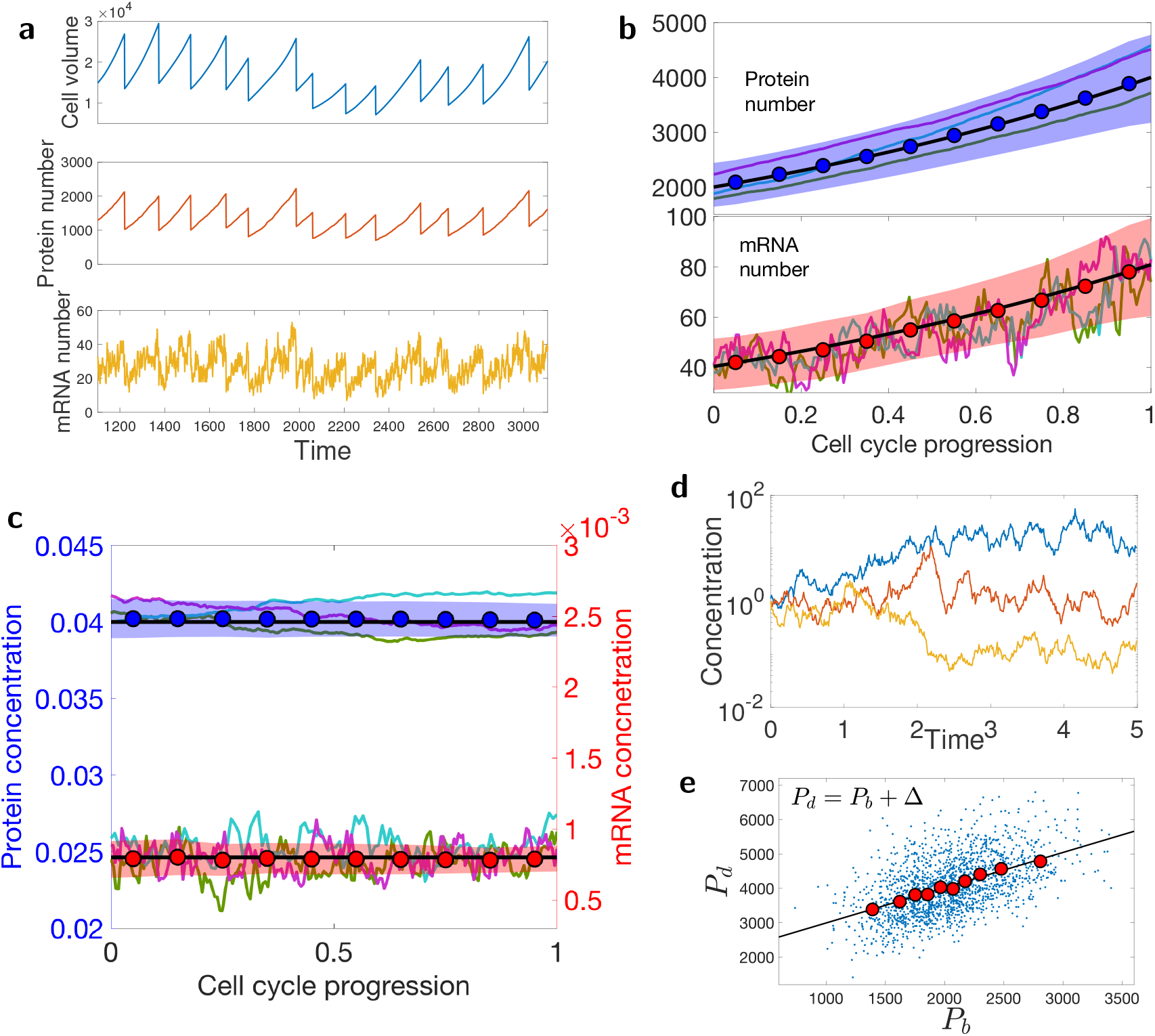
Exponential growth of the cell volume, protein number, mRNA number; the homeostasis of protein and mRNA concentrations throughout the cell cycle. (a) Numerical simulated trajectories of cell volume, protein number, and mRNA number are shown (*ϕ_i_* = 0.018). (b) The averaged values of protein and mRNA number of a highly expressed gene (*ϕ_i_* = 0.04), are shown (circles) with 3 single trajectories in the background. The black lines are theoretical predictions of Equations (4a, 4b). The average is over 130 cell cycles. The color band represents the standard deviation (same for (c)). (c) The averaged values of protein and mRNA concentrations of the same gene as in (b) are shown (circles). The black lines are theoretical predictions of Equations (5a, 5b). Three trajectories are shown in the background. (d) Three trajectories of diverging concentrations in the scenario where the protein number and cell volume grow independently. See the numerical details in Methods. (e) The scatter plot of the protein numbers at cell division (*P_d_*) *v.s.* the protein numbers at cell birth (*P_b_*). The circles are binned data. The black line is a linear fit of the binned data with slope 1.03, consistent with the adder correlations.

The corresponding trajectories of protein and mRNA concentrations are shown in Figure 2c, with bounded fluctuations around the predicted averaged values (black lines). In contrast, if the protein number and cell volume grow exponentially but independently, the ratio between them will diverge as the effects of noise accumulate, exhibiting a random walk behavior (Figure 2d). Considering the cell cycle dependence of mRNA number and the homeostasis of protein concentration throughout the cell cycle, the experimental observation in *E. coli* showing negligible correlations between mRNA number and protein concentration [47] seems to be a natural result of the cell cycle effect [48].

Within our model, we may also study the protein number dynamics: how does the protein number at cell division correlate with that at cell birth? We find that the correlations follow an “adder” (i.e. the number of new *proteins* added is uncorrelated with the number at birth), as shown in Figure 2e. While this has been quantified in various organisms with respect to cell volume [8, 9, 11, 49–51], checking correlations between protein content at cell birth and division has received significantly less attention [52, 53]. Related to this, we study the auto-correlation function of protein concentration in time. We find that the auto-correlation function is approximately exponential, with a correlation time bounded from below by the doubling time (Figure S2). Both of these results provide experimentally testable predictions.

### Effect of finite duration of gene replication

So far, we considered a constant *ϕ_i_* throughout the cell cycle assuming an instantaneous replication of the genome. In this section, we relax this condition and study the effects of finite DNA replication time. We consider the bacterial model of DNA replication, specifically, *Escherichia coli*, for which the mechanism of DNA replication is well characterized [54]. The duration of DNA replication is constant, and defined as the C period. The corresponding cell division follows after an approximately constant duration known as the D period. Details of the DNA replication model are in the Methods. In Figure 3a, we show the time trajectories of the gene allocation fraction, mRNA concentration and protein concentration of one gene for a doubling time of *T* = 30 min with *C + D* = 70 min. The DNA replication introduces a cell cycle dependent modulation of *ϕ_i_*. The abrupt increase of ϕi corresponds to the replication of the specific gene *i* (Figure 3a) *ϕ_i_* → 2*ϕ_i_*. However, as other genes are replicated, the relative fraction of gene *i* in the total genome decreases. This modulation propagates to the mRNA concentration which essentially tracks the dynamics of *ϕ_i_* due to its short lifetime. The modulation of mRNA concentration affects the protein concentration as well, yet with a much smaller amplitude. These results can be tested experimentally by monitoring the DNA replication process and mRNA concentration simultaneously. We predict a quickly increasing mRNA concentration after the gene is replicated, followed by a gradual decrease of mRNA concentration until the next round of replication.

**Figure 3.**
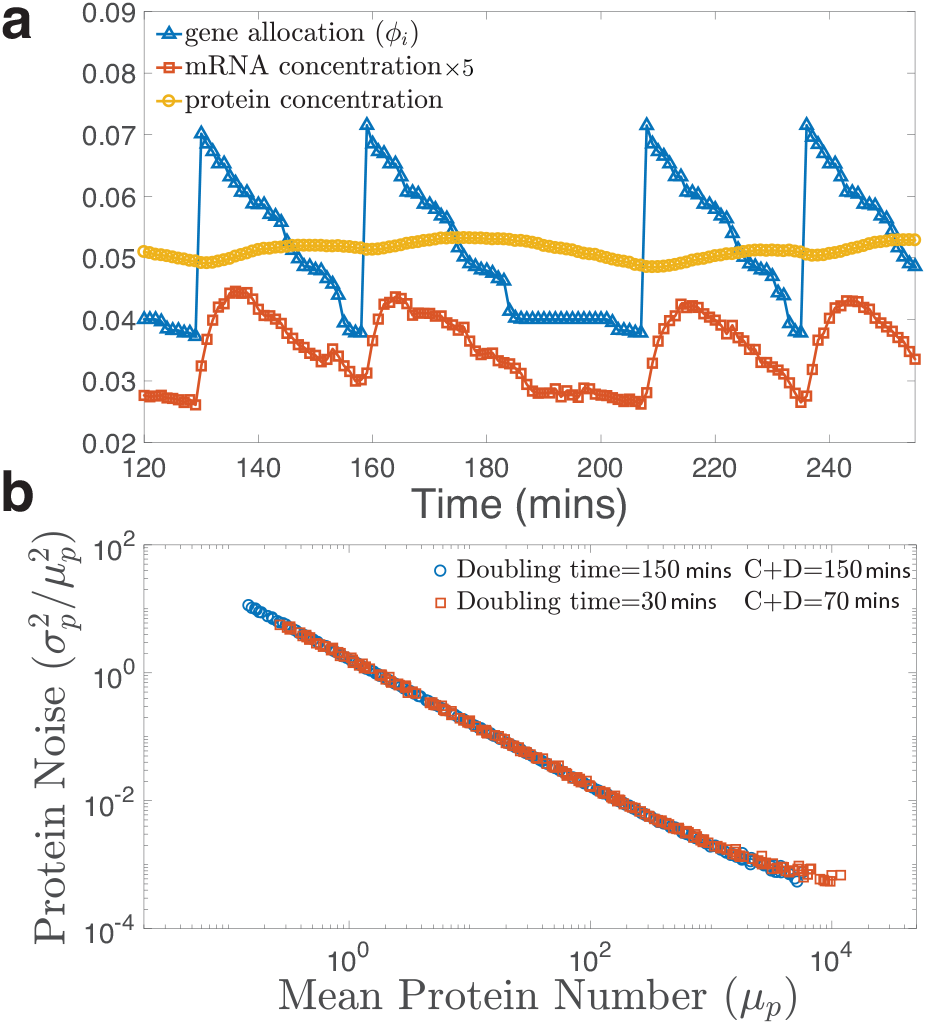
Effects of finite duration of DNA replication. (a) The time trajectory of gene allocation fraction, mRNA concentration and protein concentration of a high copy number protein (*μ_p_* ≈ 10^4^, see (b)). The doubling time *T* = 30 min, and we use the values of the C and D periods from Ref. [54], namely, *C* = 35 min and *D* = 35 min. In this situation, the cell undergoes DNA replication throughout the cell cycle. Nevertheless, the noise in *ϕ_i_* does not propagate to the noise in protein concentration significantly. The value of mRNA concentration is 5 times amplified for clarity. (b) An exponentially growing population is simulated (See Methods). The noise magnitude is quantified as the square of CV of protein concentrations. The mean protein number (*μ_p_*) is the protein number per average cell volume. Gene dosage effects due to DNA replication do not generate a significant global extrinsic noise. Two different doubling times are considered.

Noise in gene expression can be classified as intrinsic and extrinsic noise [55]. While intrinsic noise is due to the stochastic nature of the chemical reactions involved in gene expression, extrinsic noise is believed to be due to the fluctuations of external conditions and common to a subset of proteins. Experiments have revealed a global extrinsic noise that affects all protein concentrations in the genome [47, 56, 57]. Because all genes are subjected to the finite duration of DNA replication, it is tempting to attribute the finite duration of DNA replication as one of the main sources of global extrinsic noise [33]. Within our model in the previous section (constant *ϕ_i_*’s throughout the cell cycle), there is no global extrinsic noise (Figure S3). A global extrinsic noise may emerge after we introduce the time-dependent *ϕ_i_* due to DNA replication. However, we find that the coefficient of variation (CV, the ratio between standard deviation and mean) of the most highly expressed proteins is only about 0.02 within the growing cell model (Figure 3b), much smaller than that found in experiments [47, 56].

A unified phase diagram of gene expression and cellular growth

Experimental observations on *E. coli* [30] and budding yeast [31] support our assumption that ribosomes are limiting for translation. Experimental observations on plant and mammalian cells [18–20] and fission yeast [36] are also consistent with our assumption that RNA polymerase is limiting for transcription. However, as we discussed in the first section, in the same experiments on fission yeast [36] DNA became limiting for transcription at low DNA concentration. Therefore, we cannot exclude the possibility that in some cases because RNAPs are too abundant, DNA becomes the limiting resource for transcription rather than the number of RNAPs. Similarly, when ribosomes are too abundant relative to the transcript number, the limiting factor for translation becomes the transcript number rather than ribosome number.

In this section, we generalize our model by assuming that each gene has an upper bound on the number of RNAPs (*n_s_*) than can simultaneously work on it. A possible extreme case is that the gene is fully loaded with RNAPs, on which RNAPs are only constrained by steric hindrance. The same assumption is made for mRNA with an upper bound of ribosomes (*r_s_*) that can work on it simultaneously. We remark that the exact mechanism of DNA and mRNA saturation is beyond our coarse-grained model. If the number of RNAP (ribosome) is above the upper bound, the transcription (translation) rate is limited by the gene (mRNA) number, in a similar fashion to the constant rate models.

We define the protein-to-DNA ratio (PTD ratio) as the sum of protein numbers divided by the sum of effective gene numbers,

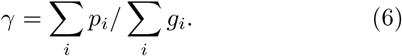

As the PTD ratio becomes larger, *e.g.*, due to a sufficiently large cell volume with a fixed number of gene, the number of RNAPs (ribosomes) will exceed the maximum load the total genes (mRNAs) can hold. We have discussed thoroughly Phase 1 (neither DNA nor mRNA is saturated) earlier and we summarize our predictions on the transition from Phase 1 to other phases in the following.

*Phase 2:* In Phase 2, the limiting factor in transcription becomes the gene copy number and the transcription rate is proportional to the gene copy number (Figure 4b). The threshold PTD ratio for the transition from Phase 1 to Phase 2 is (Methods),

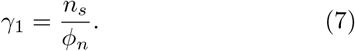

**Figure 4.**
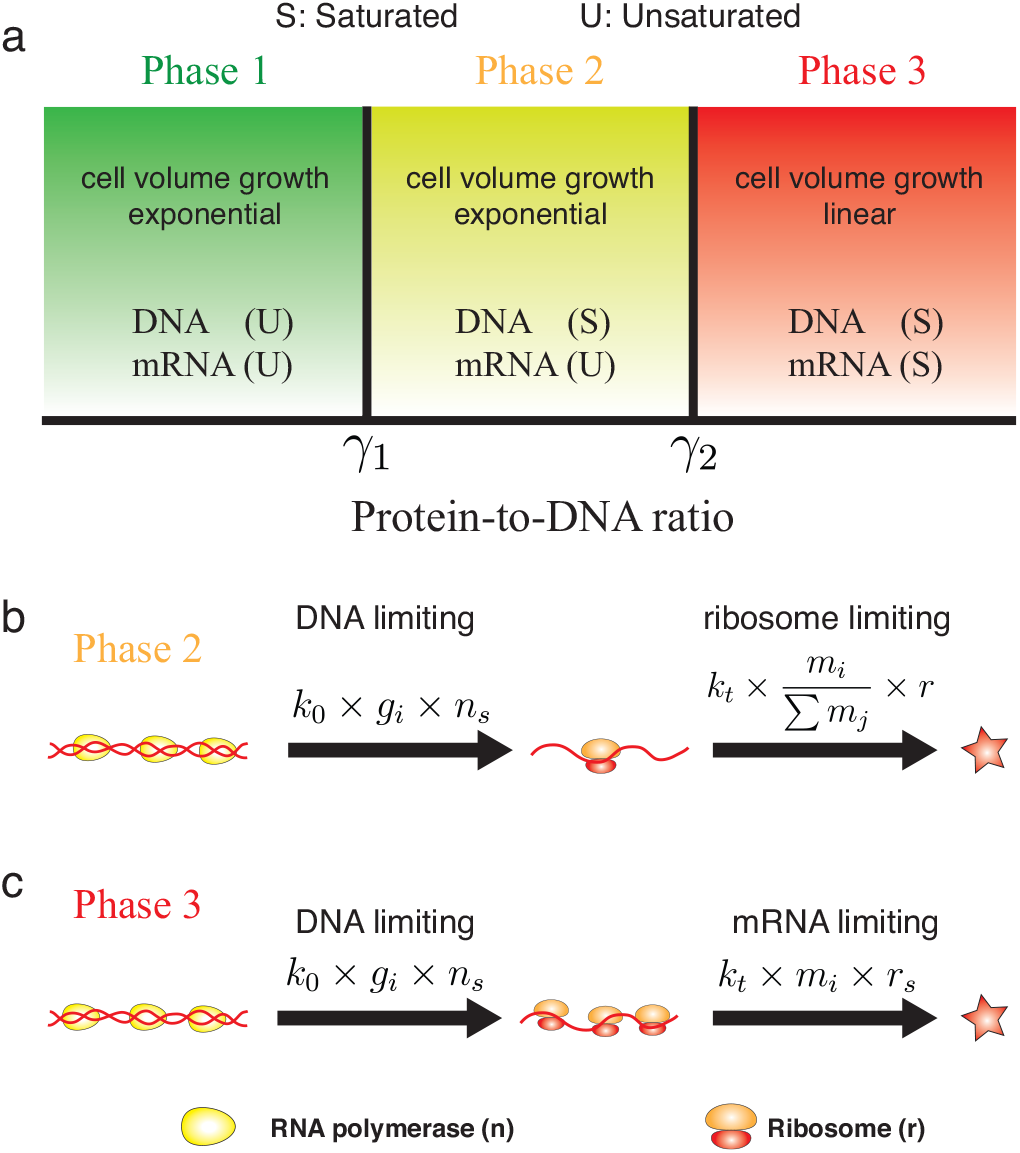
Phases of gene expression and cell volume growth. (a) Theoretical phase diagram of gene expression and cellular growth within our model. The x axis is the protein-to-DNA ratio (*γ*). When *γ* < *γ*_1_, neither DNA nor mRNA is saturated. The mRNA number, the protein number and the cell volume all grow exponentially with the growth rate set by the fraction of ribosomal gene in the total genome (*ϕ_r_*). When *γ*_1_ < *γ* < *γ*_2_, DNA is saturated but mRNA is not. The protein number and the cell volume still grow exponentially while the mRNA number is a constant proportional to the gene number. When *γ* > *γ*_2_, both DNA and mRNA are saturated. The protein number and cell volume grow linearly, and the cell volume growth rate is set by the genome copy number. (b) The gene expression dynamics in Phase 2. In this phase, DNA becomes saturated by RNAPs, therefore, the transcription rate becomes proportional to the effective gene copy number, *g_i_*. *n_s_* is the upper bound of RNAPs that can work on one gene simultaneously. The translation rate is the same as in Phase 1. To simplify the formula, we assume all ribosomes are active (to include the effect of an inactive fraction, r should be replaced by *f_a_r*). (c) The gene expression dynamics in Phase 3, in which both DNA and mRNA are saturated. The translation rate becomes proportional to the mRNA number. *r_s_* is the upper bound of ribosomes that can work on one mRNA simultaneously.

Here *n_s_* is the upper bound of the number of RNAPs that can work on one gene, *ϕ_n_* is the gene allocation fraction of RNAP. Because mRNA is not saturated, the protein number and the cell volume grow exponentially with the same growth rate as Phase 1, Eq. (3), and the homeostasis of protein concentration is still valid. However, because the production rate of mRNA is now proportional to the gene copy number, the mRNA concentration is not constant anymore as the cell volume grows and becomes inversely proportional to the protein-to-DNA ratio (Methods). We remark that in Phase 2, even though the transcription rate doubles after the genome is replicated, the translation rate is still proportional to the relative fraction of mRNA in the total pool of mRNAs. Therefore, the protein concentrations are still independent of the genome copy number. Recent proposed theoretical models of gene expression are consistent with this phase [58].

*Phase 3:* As the cell keeps growing, mRNA may get saturated as well. The limiting factor in translation is now the mRNA copy number (Figure 4c). The threshold PTD ratio for transition from Phase 2 to Phase 3 is (Methods)

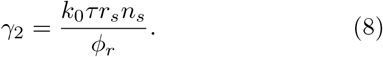

Here *r_s_* is the upper bound of the number of ribosomes that can work on one mRNA. In this phase, the translation rate is proportional to the mRNA number and the protein number grows linearly as *ṗ_i_* = *k_t_k*_0_*g_i_τn_s_r_s_*, with a linear growth rate proportional to the gene number. Therefore, within the assumption that the cell volume is dominated by the total protein number, the cell volume grows linearly as well with the linear growth rate proportional to the total gene number,

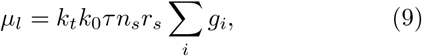

and therefore proportional to the genome copy number, *n_g_*. As in Phase 2, the mRNA concentration decreases as the cell volume grows, however, the protein concentration is still constant with the average protein concentration equal to the gene allocation fraction (〈*c_i_*〉 = *ϕ_i_*, Methods). In Phase 3, even though the cell volume grows linearly, the population still grows exponentially with a population growth rate. However, there is no general relation between the ribosomal fraction in the proteome and the population growth rate, in contrast to the growth law in Phase 1 and 2. We summarize the predicted phase diagram of cellular growth in Figure. 4.

To gain some sense regarding the parameters associated with our proposed phase diagram, we estimate the PTD ratio of *E. coli*. Considering the typical cell volume of *E. coli* as 1 *μm*^3^, the protein density as 3 × 10^6^ proteins/*μm*^3^ and the total number of protein-coding genes in *E. coli* as 4000 [59], we estimate the protein-to-DNA ratio for *E. coli* as γ ~ 1000. Estimates of the two threshold values of PTD ratios (see Methods) suggest that *γ*_1_ ~ 1500 and *γ*_2_ ~ 20000.

We find that *E. coli* cells are typically in Phase 1, but not too far from Phase 2. We remark that the actual threshold values of PTD ratio for the transitions between different growth phases may be affected by other factors, e.g., the heterogeneous size of genes, but we propose that the general scenario of the transition from Phase 1 to Phase 3 as the protein-to-DNA ration increases should be generally applicable. As the PTD ratio increases, we predict a transition from exponential growth to linear growth for protein number and cell volume. We propose future experiments to study the potential transition from exponential to linear growth of cell volume, for example using filamentous *E. coli* by inhibiting cell division and gene replication. Similar experiments can also be done for larger cells, e.g., mammalian cells, in which the transition from exponential growth to linear growth of cell volume may be easier to achieve. Preliminary results from experiments measuring the growth of cell mass of mammalian cells by inhibiting cell division indeed show a crossover from exponential growth to linear growth when the cell mass is above a threshold value [60], consistent with our prediction.

## CONCLUSION

In this work, we propose a coarse-grained model of stochastic gene expression incorporating cell volume growth and cell division. In the first part, we consider the biological scenario that RNAPs are limiting for transcription and ribosomes are limiting for translation. In other words, neither DNA nor mRNA is saturated. We find that the limiting nature of ribosomes in the translation process leads to the exponential growth of protein numbers. The limiting nature of RNA polymerase and its exponential growth lead to the exponential growth of mRNA numbers. Homeostasis of protein concentrations originates from the fact that ribosomes make all proteins. Homeostasis of mRNA concentration comes from the resulting bounded concentration of RNAPs. Our model is consistent with the constancy of mRNA and protein concentration as the genome copy number varies since the transcription rate depends on the relative fraction of genes in the genome rather than its absolute number [22].

During DNA replication, we find that the gene allocation fraction *ϕ_i_* for one specific gene doubles after the gene is replicated but decreases afterwards since other genes are replicated as well and compete for RNAPs. This prediction can be tested by monitoring the mRNA concentration and the copy number of one gene throughout the cell cycle. Furthermore, we extend our model to more general cases in which DNA and mRNA can be saturated by an excess of RNAP and ribosome. We find three possible phases of cellular growth as the protein-to-DNA ratio *γ* increases. A transition from exponential growth to linear growth of protein number and cell volume is predicted. In the future, it will be interesting to study the interplay between the global interactions which are the focus of this work and local interactions between genes. Our model provides an alternative model to constant rate models to study genetic networks, which would be advantageous when cell cycle effects are important. Another potential extension of our model is to include metabolic proteins and investigate the effects of nutrient limitation on the gene expression and cell volume growth.

## METHODS

### Derivation of protein and mRNA concentrations

We define the fraction of mRNA *i* in the total mRNA pool as *f_i_* = *m_i_*/Σ*_j_ m_j_*, and the concentration of mRNA and protein of gene i as *cm_i_* = *m_i_*/*V*, *c_i_* = *p_i_*/*V*. We denote the RNAP and ribosome concentration as c*n* and *c_r_*. According to Equations (1a–1c), the deterministic equations of the above variables then become

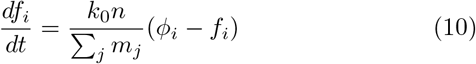

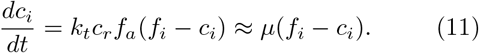

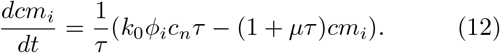

Using the condition that mRNA degradation time is much smaller than the doubling time (*μτ* ≪ 1), we find the fixed points for the dynamics of *f_i_*, *c_i_*, and *cm_i_*. These are, respectively, *ϕ_i_*, *f_i_*, and *k*_0_*ϕ_i_c_n_τ*. Replacing *f_i_* by *ϕ_i_* and *c_n_* by *ϕ_n_*, we obtain the approximate version of the above equations, Eq. (5a,5b).

### Simulations of independent growth model

In the growth model corresponding to Figure 2d, we assume the protein number and cell volume grow exponentially and independently,

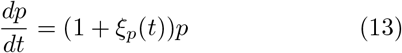

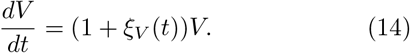

Here, *ξ_p_*(*t*), *ξ_v_*(*t*) are white noise terms, with the autocorrelation function, 〈*ξ_p,v_*(0)*ξ_p,v_*(*t*)〉 = *A_p,v_δ*(*t*). In Figure 2d of the main text, we choose *A_p_* = *A_v_* = 1.

### Simulations of growing cell model

We simulated Equations (1a,1b,1c), fixing *r_b_*, *n_b_*, *b*_0_, *ϕ_r_, f_a_, I_c_, τ* as well as the growth rate *μ*. Other parameters are inferred given the above values, *e.g*., *ϕn* = *n*_6_*ϕ_r_/r*0, *k_t_* = *μ*/(*ϕ_r_f_a_*), *k*_0_ = *k_t_f_a_r_b_*/(*b*_0_*n_b_*). We fix the time step δt so that the probability for one event to happen during a time step is smaller than 0.1. We track one of the daughter cells after cell division.

### Gene dosage effects

In reality, the gene allocation fraction *ϕ_i_* changes during the cell cycle due to the finite duration of DNA replication. In this section we introduce the modified version of the gene expression model incorporating DNA replication. Although our model is general, we focus on DNA replication in bacteria for concreteness, specifically *E. coli* where this process is very well characterized. We expect our conclusions to be generally valid. Furthermore, we refine our model for cell division, assuming that the initiator protein triggers the initiation of DNA replication rather than cell division, with the threshold *I_c_* proportional to the number of origins of replication [54, 61] (the number of which doubles at each initiation). We assume that the cell division takes place a fixed time *C + D* after initiation of the DNA replication, where *C, D* are respectively the time for DNA replication and the time between the completion of DNA replication and cell division. The number of origins reduce by half at each cell division. Other details are the same as in the main text. Each gene doubles its copy number during the *C* period, and we choose this gene replication time to be randomly and uniformly distributed across all genes. When a gene *i* replicates,

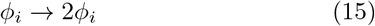

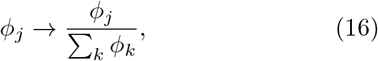

where the second equation accounts for the normalization of the gene allocation fraction. We choose the experimentally reported *C* and *D* and cell doubling time from Ref. [54]. In Figure 3a, we simulate the model by tracking one daughter cells. In Figure 3b, we track all the cells in an exponentially growing population, which starts from 100 cells to 5000 cells.

### Simulations of gene activation

We generalize the constitutive expressed genes considered in the main text to include a single regulated gene by considering a random telegraph process of the effective gene copy number [1],

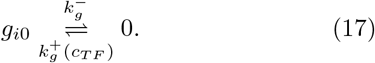

Here the gene deactivation rate 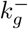 is constant, and the activation rate is set by the concentration of transcription factor through positive regulation, 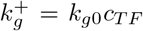. Here, *k_g_*0 is constant. When gene *i* is active, the corresponding gene allocation fraction follows *ϕ_i_* = *g*_*i*0_/Σ*_j_ g_j_*, and when it becomes deactivated *ϕ_i_* = 0. Note that here we only consider one regulated gene *i*, but the changing gene allocation of gene *i* also affects other genes’ allocation fraction. We simulate the model in Phase 1, and the deactivation of gene i increases other genes’ allocation fraction as *ϕ_j_* → *ϕ_j_*/(1 – *ϕ_i_*), with *ϕ_i_* = *g*_*i*0_/Σ_*j*_ *g_j_*.

Simulated trajectories of gene allocation fraction, mRNA number, protein number and cell volume are shown in Figure S1.

### General model of gene expression

We consider the generalized equation of mRNA number, Eq. (1a) in the deterministic limit as

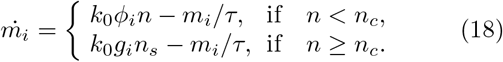

Here *n_c_* is the threshold number of RNAPs above which DNA starts to be saturated, in which case the transcription rate becomes proportional to the effective gene copy number *g_i_* and independent of the RNAP number. For one gene, the maximum load of RNAP that it can hold is *g_i_n_s_*, where *n_s_* is the maximum number of RNAPs that a single copy of constitutively expressed gene (*g_i_* = 1) can hold. *n_c_* can be computed as

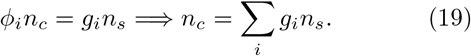

We also generalize the growth of protein number from Eq. (1c) to

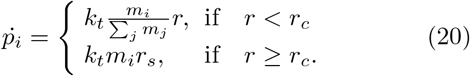

Here *r_c_* is the maximum number of ribosomes above which mRNA starts to be saturated. We drop the fraction of actively working ribosomes since it is often a constant depending on the growth condition [30]. *r_s_* is the maximum number of ribosomes one mRNA can hold. We can calculate *r_c_* as

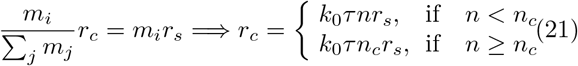

Given Eqs. (18, 20), we obtain four possible phases: (i) *n < n_c_, r < r_c_*, (ii) *n > n_c_, r < r_c_*, (iii) *n > n_c_, r > r_c_*, and (iv) *n < n_c_, r > r_c_*. Given a fixed value of *ϕ_r_* and *ϕ_n_*, either (ii) or (iv) is possible. Realization of (ii) requires that *n* > Σ_*i*_ *g_i,n_s__* and *r < k*_0_*ττ_s_ g_i_n_s_*, therefore

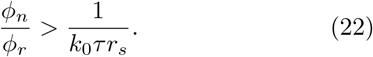

In cases where Eq. (22) breaks down, a finite fraction of ribosomes are not utilized. Based on various recent works [30, 32], this would be highly inefficient for cellular growth and we expect Eq. (22) to hold for fast proliferating cells. This requires a large fraction of genes in the genome making ribosomes that can not work on translating because mRNAs are saturated. Since ribosomes are typically more expensive to make than other proteins, we assume the biological scenario, Eq. (22) will be satisfied.

From Eq. (19) and using *n*/Σ*_i_ p_i_ = ϕ_n_*, we obtain the threshold PTD ratio for the transition from Phase 1 to Phase 2,

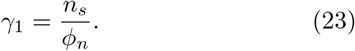

In Phase 2, the average mRNA concentration becomes

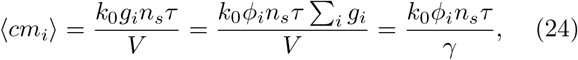

which is inversely proportional to the protein-to-DNA ratio.

From Eq. (21) and using *r*/Σ*_i_ p_i_ = ϕ_r_*, we obtain the transition PTD ratio from Phase 2 to Phase 3 as,

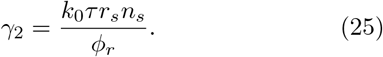

In Phase 3, the mRNA concentration is the same as Phase 2. Because the protein number grows linearly and the cell volume is the sum of all proteins, the protein concentration is the same as Phase 2 and Phase 1, 〈*c_i_*〉 *g_i_*/Σ*_i_ g_i_ ϕ_i_*.

### Estimation of the threshold protein-to-DNA ratios for *E. coli*

We approximate the upper bound of RNAP number working on a single gene as roughly equal to the number of RNAPs on a typical gene (~ 10^3^ base pairs) when half of the gene is occupied. The linear size of RNAP is about 5 *nm*, and the length of one base pair is about 0.3 *nm*, leading to the estimate *n_s_* ~ 30. A similar calculation for the upper bound of ribosome on a single mRNA leads to *r_s_* ~ 10 since ribosome’s linear size is about 3 times ı larger than RNAP [59].

We take *ϕ_r_* ≈ 0.2 according to the Ref. [30], and estimate the gene allocation fraction of RNAP to be *ϕ_n_* ~ 0.02 since the number of RNAPs in *E. coli* is roughly 10% of the number of ribosomes [59]. We estimate the life time of mRNA as 5 mins [59].

We estimate the transcription rate of one RNAP by considering two potential limiting steps in transcription and take the slower one. First, assuming the initiation 0 of transcription is diffusion limited, we could estimate ı the time scale for one RNAP to bind the transcription site as Δ*t* ~ 1*μm*^2^/(0.2*μm*^2^/*s*) ~ 5*s* using the measured diffusion constant of RNAP [62, 63]. Second, we could also estimate the elongation time as the typical length of gene divided by the elongation rate of RNAP, Δ*t* ~ 1000*nt*/50(*nt/s*) ~ 20*s* [59]. Taking the slower time scale from the above two calculations, we estimate *k*_0_ ≈ 0.05*s*^−1^. Finally, we compute *γ*_1_ and *γ*_2_ using the above estimated parameters, and obtain *γ*_1_ ~ 1500, *γ*_2_ ~ 20000.

## AUTHOR CONTRIBUTIONS

All authors conceived the work, carried out the work, and jointly wrote the manuscript.

## ACKNOWLEDGMENTSS

We thank Naama Barkai, Ido Golding, Andreas Hilfinger, Po-Yi Ho, Andrew Murray, Johan Paulsson, Leonardo A. Sepuúlveda, and Sven van Teeffelen for useful discussions related to this work. AA thanks the A.P. Sloan foundation, the Milton Fund, the Volkswagen Foundation and Harvard Deans Competitive Fund for Promising Scholarship for their support. JL was supported by the George F. Carrier fellowship and the National Science Foundation through the Harvard Materials Research Science and Engineering Center (DMR-1420570)

